# RAD-Behavior (Recombining Atomized, Discretized, Behavior): A new framework for the quantitative analysis of behavioral execution

**DOI:** 10.1101/739151

**Authors:** Russell A. Ligon, Edwin Scholes, Michael J. Sheehan

## Abstract

The ability to precisely describe and numerically evaluate organismal phenotypes is a prerequisite for addressing most questions in evolutionary biology and ecology. The quantification and comparison of behavior, loosely defined as an external response to stimuli, is particularly challenging because the myriad axes of variation that exist make comparisons, both within and among species, difficult. Such evaluations often boil down to comparisons of time-budgets (e.g. relative investment in courtship displays) or probabilities (e.g. likelihood of engaging in a class of behaviors in a particular context) – which we refer to as *behavioral strategies*. A focus on variation in behavioral strategies underlies most research in evolutionary and ecological studies of behavior. Equally important, however, is perhaps the question of ‘how’ animals are actually performing the complex motor sequences that comprise behaviors (i.e. *behavioral execution*). What are the patterns of movement, the relative transition rates, and kinematics underlying the behaviors exhibited in particular contexts? Understanding how behavioral execution differs among individuals, populations, and species has the potential to provide new insights into the factors shaping variation in behavior and the processes shaping behavioral evolution at different scales. Here, we propose a broad framework for comparing behavioral execution (RAD-behavior: recombining atomized, discretized behavior) that leverages string-matching/bioinformatic tools to understand phenotypic variation in behavioral execution and which holds the potential to yield novel insights about the evolutionary ecology of behavior at multiple scales.

---

Computationally intensive, data-rich approaches have facilitated new biological insights at many levels of analysis in recent years. However, the ability to leverage vast quantities of data into meaningful answers to longstanding biological questions depends on the development and implementation of appropriate analytical toolkits. Here, we propose a framework for measuring and comparing behavior (RAD-behavior: recombining atomized, discretized behavior) that opens the door to bioinformatic analyses of behavior. Many rigorous and computationally efficient tools exist to tackle questions about variation in, and the evolution of, genetic sequence information. RAD-behavior facilitates similar analyses for behavior, enabling the numeric comparison of behavioral sequences, allowing the comparison of behavioral sequences across taxa, enabling efficient searches of large behavioral databases, and providing an important advance in our attempts to better understand phenotypic variation in behavioral execution. Collectively, this approach has broad potential to yield novel insights about the evolutionary ecology of behavior at multiple scales.

Though several high-throughput behavioral classification techniques have recently been developed^1–7^, questions remain about how to leverage the resulting datasets generated from such techniques^8–10^. In this paper, we describe an analytical pipeline for behavioral analyses to answer questions about behavioral homology, repeatability, innovation, and performance by leveraging the varied and flexible toolkit commonly employed by genomic researchers. Collectively, we term this analytical and conceptual framework *RAD-behavior (Recombining Atomized, Discretized behavior)*. The basic idea of RAD-behavior is simple: break down behavior into subunits (‘atoms’) that are shared across individuals and/or species, assign discrete character states for each behavioral atom at every time point, then recombine these discrete character states to create strings that collectively describe behavioral execution. This approach is consistent with, and an extension of, the idea that complex behaviors can be “created from the flexible combination of a small set of modules”^11^, as well as the idea that many behaviors can be described as sequences of physical orientations/postures^9^. The RAD-behavior framework embraces the long history of homology thinking in behavioral research^12–14^, has its roots in classical ethology^15^, and builds on a number of creative approaches developed for the description and analysis of movement^16,17^.

Here, we describe the RAD-behavior framework in sufficient detail to enable interested researchers to incorporate this framework into their particular study systems and research programs. Specifically, we discuss how, with a rigorous behavioral atomization and discretization pipeline, bioinformatic tools like BLAST (Basic Local Alignment Search Tool)^18^, MEME Suite (Multiple EM for Motif Elicitation)^19^, and MAFFT (Multiple Alignment using Fast Fourier Transforms)^20^ can be used to find putatively-homologous behavioral sequences (within and across individuals, and even across species), find and rank repetitive behavioral elements, identify behavioral modules shared across species, and pinpoint evolutionary innovations among divergent taxa. The quantitative measures of behavior that this analytical pipeline offer have the potential to yield new insights into ecological and evolutionary drivers of behavioral diversification, and we illustrate some of these possibilities with exemplar demonstrations from a comparative dataset of courtship behaviors from the birds-of-paradise. However, we emphasize that RAD-behavior is neither a tracking tool nor a computational program. Rather, RAD-behavior is a conceptual framework that can incorporate diverse data types (e.g. from manually scored videos or automated body-region tracking of targets) to facilitate the creation of detailed strings of behavioral data that can be analyzed using powerful computational and bioinformatic tools to answer new questions, old and previously unanswerable questions, and questions that have yet to be asked about behavioral variation across scales.

## Challenges of Comparing Behavior

> ”…the comparison of many ethograms is a nontrivial statistics problem…”^21^

Quantitative comparisons of behaviors are challenging for many reasons^10^. Key among these challenges is the difficulty in scoring sets of behaviors that may share few obvious similarities. What behavioral elements should be scored and compared? The challenge of “identifying the most meaningful and useful dimensions and descriptive units of behavior is…*the* problem”^22^ of comparative behavioral research. Items that are highly divergent are difficult to compare, like the clichéd “apples vs. oranges” comparison used to illustrate the non-comparability of two or more objects. How does one compare the greenness of an apple to the orangeness of an orange? Do we compare the number of carpels? Do we compare the symmetry? The difficulty of making meaningful comparisons across things that are really quite different is in no way unique to the evolutionary study of behavior, yet this difficult challenge has hampered our understanding of behavioral evolution. Molecular geneticists and population genomicists get around *part* of this issue by drilling-down as deeply as biologically meaningful – getting at the raw genetic sequence variation among individuals, populations, and species. Their answer can be used as a guide – remove higher-order complexity until the underlying units facilitating differentiation are comparable. Said another way, evolutionary biologists interested in comparative studies of animal behavior can break down complex behaviors into sub-units that are shared across individuals and species. Going back to our fruit analogy, apples and oranges are almost incomparable if we simply take a giant bite of each (peel included), but if we identify the proteins and sugars in each, we may be able to make quantitative comparisons that facilitate new insights into the key factors that differentiate these classically incomparable fruits.

A second significant challenge in the study of behavior lies in the temporal alignment, or misalignment, of the behaviors being performed. Unlike the genetic sequence from a given chromosome which has a ‘start’ and ‘end’, living animals are always behaving. How one defines the onset and cessation of behavioral sequences to compare among individuals and species is therefore an important and largely unresolved question. Additionally, what is the relative importance of the specific timing of a particular behavior? Does timing matter more, or less, than the order in which the overall behavioral sequences progresses? How many steps in a sequence are worth considering when analyzing behavioral patterns^23^? These are longstanding challenges in the scientific study of behavior that have been addressed in myriad and creative ways by scientists from many disciplines, frequently in ways unique to specific research aims and questions. Yet these challenges remain largely unsolved at broad scales and novel conceptual approaches are required to leverage the rapidly developing technological capabilities available to behaviorists^10^. The simplicity of RAD-behavior (breaking down complex behaviors to their core ‘**atoms**’ (Box 1), measuring and discretizing these subunits, and recombining these character states into a single string) coupled with its integrability with new technologies (e.g. machine-learning video tracking) make it well-suited as a tool for addressing longstanding questions in behavior.

### Box 1 Key Terminology

#### Behavioral Atom

Building block of behavior, conceptually aligned to the building blocks of chemistry. Just as atoms of different elements can be combined into composite molecules, different behavioral atoms can be combined by organisms into complex, composite behaviors.

#### BLAST (Basic local alignment search tool)

Search algorithm designed and optimized for searching biological sequences (e.g. DNA sequences, protein sequences). This tool allows researchers to quantitatively identify the degree to which a sequence of interest matches other sequences within a reference database.

#### Consensus sequence

A sequence representing the most frequent behavioral states at each temporally-encoded behavioral subunit site, generated following a multiple sequence alignment (MSA) from multiple execution strings.

#### Ethome

Just as a genome is a comprehensive description of the genetic material of an organism, an ethome is a comprehensive description of the behavioral states of an organism.

#### Instantaneous body configuration

Comprehensive numerical description of the organism’s body position in space (including both relative measurements between body regions and absolute measurements with respect to the vertical plane). Consecutive ‘instantaneous body configurations’ comprehensively describe an organism’s behavioral execution.

#### Principal Component Analysis

A data-reduction and transformation approach to understand broad patterns in large datasets with many variables.

#### Reference sequence

A behavioral sequence (in string form) against which comparisons of new sequences can be made and interpreted. In the case

## What constitutes a useful behavioral atom?

> “When considering anatomy, we use the notion of homology across taxa to substantiate the universality of particular forms. A future behavioral science might complete Lorenz’s vision of establishing homology in ethology.^18^”

When defining discrete behavioral subunits for use in comparative analyses, there are several important considerations. First, the complete set of behavioral subunits must be combinatorial in such a way that the combined use and scoring of subunits must enable comprehensive scoring and categorization of every behavior exhibited within the set of contexts being analyzed, across species. Second, subunits should be defined broadly enough that individual behavioral subunits should be shared across the individuals, populations, and/or species under study. If behavioral subunits are exclusive to single species, their inclusion in the overall, comprehensive ethogram should be well-justified. Third, when generating the composite list and corresponding definitions for a comprehensive, multi-species ethogram composed of behavioral subunits, the question of whether the subunits need be homologous should be considered carefully. On one hand, behavioral sequences generated using a comprehensive ethogram containing *only* homologous behavioral subunits will have the highest degree of conceptual overlap with genomic sequences, better facilitating broad, evolutionarily-influenced interpretations about evolutionary trajectories, inflection points, and behavioral modules. On the other hand, researchers may have good reason to classify certain behaviors in the same way, even if they are produced via non-homologous actions. For example, investigating the evolution of courtship behavior in birds-of-paradise from the perspective of females^24^, we might argue that shape-shifting (i.e. when individuals assume a nonbird-like shape through feather accentuation or unusual body positioning in space) is an important, relevant behavioral category that is not completely captured with additional behavioral subunits (e.g. ornamental feather accentuation). This decision should depend on the goal of the research, where such an approach would not make much sense if the goal was to understand the neural circuitry underlying the expression of particular behavior (given that shape-shifting might arise via the use of different muscles in different species) but might be justifiable when exploring the evolutionary underpinnings of behavioral complexity influencing female choice.

How do behavioral subunits differ within a larger ontology of behavior^25^? Is it reasonable to assign equal weight to a behavioral state describing the erection of a single feather and one describing the movement status of the whole organism? These behavioral atoms clearly differ in the amount of coordination and the number of muscle groups they involve. Does it make sense to restrict behavioral subunits to those that involve only equal levels of muscle engagement and coordination? Arguably, it is too early to say much about how, when, and whether different behavioral subunits should, and should not be assigned the same salience within the RAD-behavior framework. Only time and data will tell, as tests of evolutionary patterns of behavioral sequences implementing this framework are uniquely suited to evaluate how and whether macroscale patterns of evolutionary rates differ among behavioral subunits with different positions within a larger behavioral ontology hierarchy. Put another way, the RAD-behavior framework opens up novel lines of inquiry for the investigation of the processes governing behavioral evolution. One could imagine that, with sufficient justification, a weighting scheme could be applied such that particular behavioral atoms have higher weights (i.e. are represented by longer character strings within the composite string describing an organism’s instantaneous behavioral state) than others based on simulation studies and comparisons of evolutionary rates (e.g. using recently developed methodologies^26^). The flexibility of the overarching RAD-behavior analytical framework makes it well-suited to these case-by-case designations that will depend on the questions being asked by researchers.

## Recombining Atomized, Discretized Behavior(RAD-behavior): An overview

### Collecting behavioral recordings

As with many approaches for the fine-scale quantification of behavior, the RAD-behavioral approach typically relies upon video recordings of animals in contexts of interest (Figure 1a). For high-throughput analyses, individuals could be filmed simultaneously with multiple, high-definition video cameras that have been calibrated to facilitate 3-dimensional measurements of points within the frame of the cameras^27^. In certain circumstances, such an approach may not be necessary or possible, and video-recordings of subjects from a single vantage point can still be used for RAD-behavior analyses. At present, these single-viewpoint videos often, but not always^23,28^, require labor-intensive manual scoring of behaviors. The videos capturing behavioral sequences provide the basis for the subsequent steps of the RAD-behavior pipeline.

**Figure 1:**
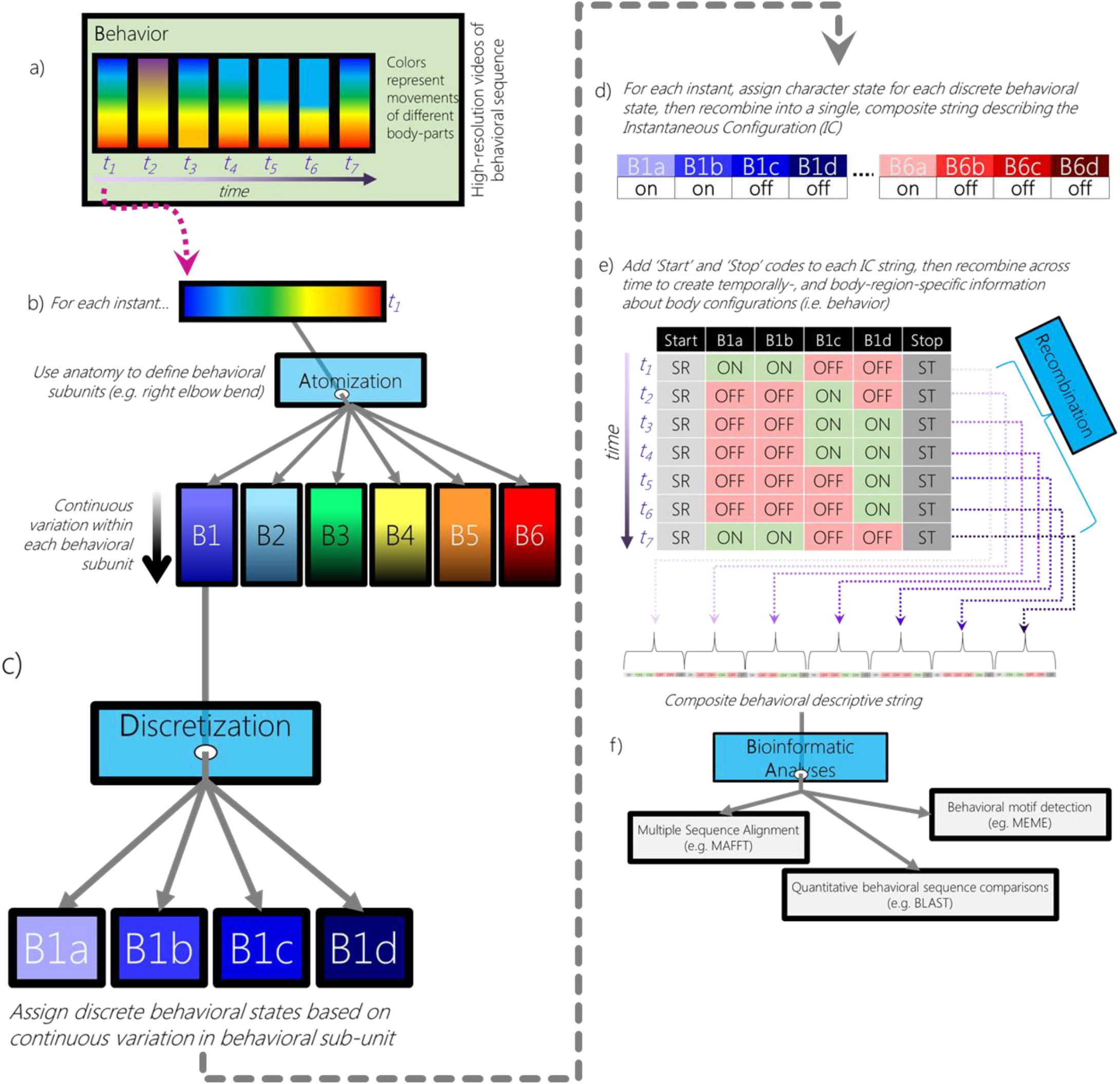
Conceptual overview of RAD-behavior analytical pipeline. Behavioral execution can be conceptualized as a series of sequential states **(a)**, where the *instantaneous body configuration* can be measured at each time point (e.g. t1-t7). The instantaneous configuration of body parts, and/or the instantaneous behavioral state can be broken down into smaller subunits (atoms), which collectively describe the organism’s state at each instant **(b)**. Each behavioral atom can be broken down into smaller, discrete states **(c)** that can be evaluated at every time point. The collective behavioral states and/or body configuration of the organism can thus be described as a character string at every analyzed time point **(d)**. Following the generation of a character string describing the instantaneous body configuration/behavioral state of the animal **(d)**, consecutive instances of these strings can be combined **(e)** to generate a composite string that collectively describes the sequences of movements and/or behavioral states of the organism. Lastly, these strings can be evaluated with a suite of pre-existing bioinformatic tools, or with custom string matching/searching algorithms **(f)**.

### Atomization

Following video capture of the behavioral sequence of interest, researchers must decide on a set of behavioral subunits, or atoms (Box 1), that collectively enable the comprehensive description of all behavioral states of all individuals in the study. The atomized ethogram must include behavioral subunits that are shared across many (if not all) individuals in the study group (including across species, for comparative studies) and these subunits should enable combinatorial pluripotency – the ability to collectively and completely describe any behavior of any organism in the study. Thus, at every time interval, a behavioral state can be determined that is the composite of all ‘active’ behavioral atoms at that instant. In cases where rigorous human quantification of behavior is the appropriate approach, behavioral atoms should arguably be based on movement patterns of body parts shared by all individuals under study. Alternatively, if the behaviors in question are used as signals, behavioral atoms that are arguably perceived (rather than produced) in the same way might constitute more informative behavioral subunits^24^. In cases where users desire a high-throughput pipeline for generating composite behavioral strings, automated tracking of body parts over space and time can be used to provide continuous measures of behavioral atoms. Specifically, focal body regions can be thought of as a ‘behavioral atoms’ that are tracked through space and time, and these temporally-specific positional data can be used to generate frame-by-frame measures of absolute (Figure 2a) and relative (Figure 2b) geometic orientation data. Currently, there are at least two readily implementable pipelines for such tracking that do not rely on the use of attached markers, both incorporating deep learning approaches to facilitate body-region identification and tracking^29,30^. As new tracking techniques and methods are developed, they can be incorporated into the RAD-behavior framework – all that RAD-behavior requires is information on the spatial arrangement of body parts over time.

**Figure 2.**
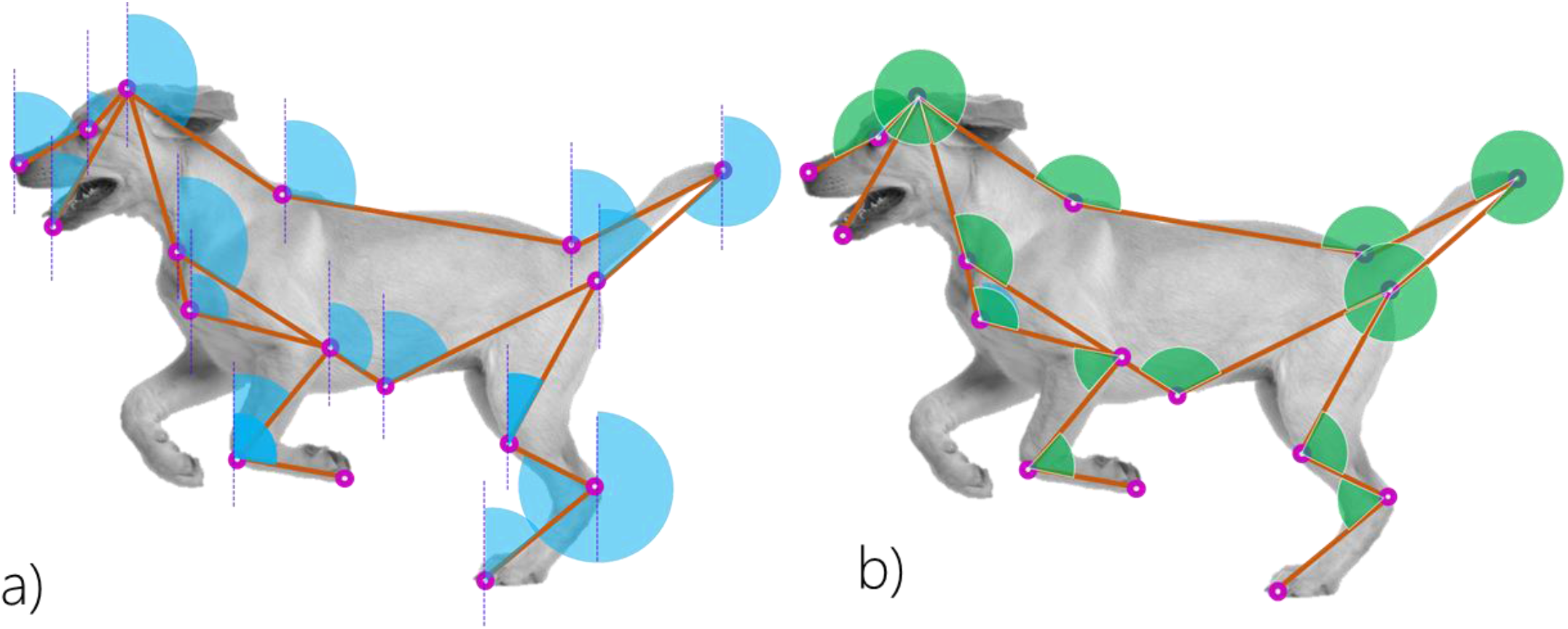
Representation of (a) absolute and (b) relative geometric orientation data which can be combined to generate an ethomic sequence using RAD-behavior.

### Discretization

Once ‘behavioral atoms’ are measured (i.e. once pre-determined body-regions are tracked through space and time, and geometric orientation information is obtained from them), a **discretization** step is implemented. Discretization is achieved by assigning threshold categories for the originally continuous numerical data describing the geometric orientation of each body region/behavioral atom (Figure 2). Thresholds should be optimized in a context-specific manner to determine optimal granularity; however, thresholds need to be identical for each behavioral sequence in a given dataset in order to facilitate comparisons between behavioral sequences. For example, when measuring the geometric configuration of body-regions, a 10° threshold may be used which effectively bins behavioral atoms into 10° units, each of which can then be classified as ‘on’ or ‘off’ at a given instant (Figure 1d). These thresholds turn the continuous measurements of the ‘behavioral atoms’ (Figure 1b) into the discrete units (Figure 1c) required for composite string creation (Figure 1d). As a result of the measurement of behavioral atoms and the discretization thresholds chosen by the user, the **instantaneous body configuration** (a comprehensive description of how the body is configured at a given moment; Box 1) of the focal individual can be encapsulated as a string of “off” and “on” values which correspond to, and provide information about, how a body is arranged in space at that instant (Figure 1d).

### Recombining discretized behavioral data into sequence data

It follows that if an individual’s instantaneous body configuration can be determined and encapsulated in the form of a string of “off” and “on” values (where each position in a sequence has only one of these ‘states’), then a series of instantaneous body configurations will necessarily describe patterns of movement over time – that is, behavior (Figure 1e). Specifically, strings detailing instantaneous body configurations can be arranged sequentially, thus creating a new composite string that captures information about how an individual’s body moves over time (Figure 1e).

After generating high-resolution, discretized behavioral strings (Figure 1d) for each time point, the data need to be converted into a sequence that can be compared across individuals, populations, and species (Figure 1e). One approach to facilitate this transformation relies on using a behavior mapping key (Figure 3a) to generate a large behavior-by-time matrix where each behavioral atom is assigned a specific location (i.e. column) within the matrix, and rows represent discrete time-intervals (Figure 3b,c). To ensure uniformity of behavioral subunit position across a given study, a behavioral subunit-position mapping key (Figure 3a) links each behavioral subunit to a specific location in the sequences to be generated, facilitating all subsequent comparisons. Once this mapping is completed, it cannot be changed unless all behavioral sequences are re-converted to RAD-behavior strings using the new mapping scheme. Depending on the sequence matching tools and alignment approaches used, the connection between the specific behavioral subunit and its string position can be i) randomized (in cases where string matching algorithms are focused on overall sequence similarity irrespective of consecutive chains of matches) or ii) ordered by body region (if string searching algorithms reward ‘bonuses’ for consecutive matches).

**Figure 3.**
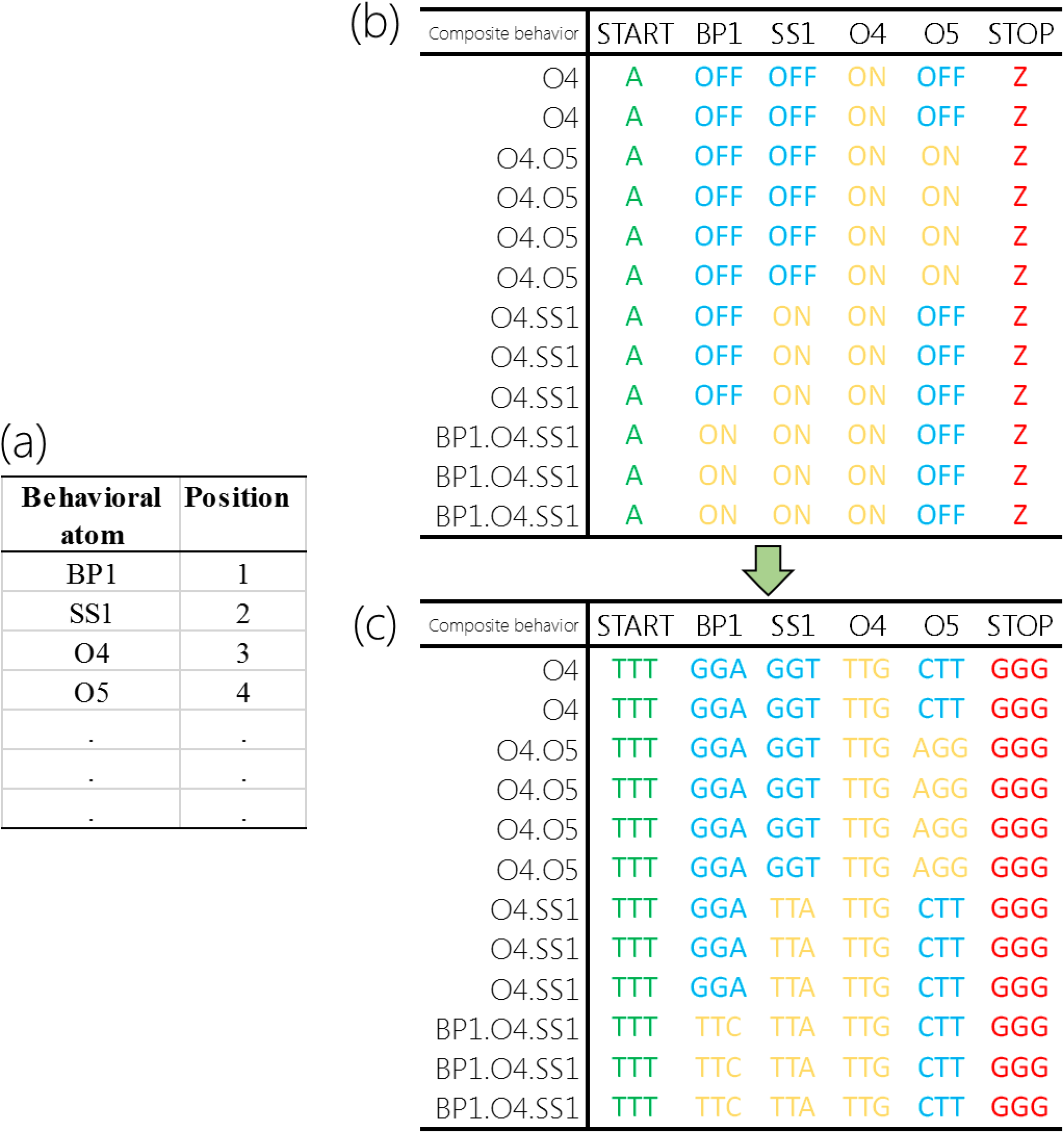
Visualization of (a) behavior-position mapping key and (b) Behavior x Time (BT) matrices used for sequence creation. The Behavior-Position mapping key (a) links each behavioral atom to a specific location within the sequences to be generated, facilitating all subsequent comparisons. The BT matrices (b,c) have N rows, corresponding to the number of measurements obtained across time, and M columns corresponding to the number of behavioral atoms scored plus 2 (for ‘start’ and ‘stop’ codes). The behavior-position key will dictate the which columns in the BT matrcies correspond to which behavioral subunit. After each discretized behavioral subunit is scored as 0 (‘OFF’) or 1 (‘ON’) at each of N timepoints (b), these states can be converted to character strings that are unique to the behavioral subunit and state.

After generating a behavior mapping key, the N x M behavior-by-time matrix is created (Figure 3b), where the N rows correspond to the number of measurements obtained, ordered sequentially, and M columns correspond to the number of behavioral subunits scored plus 2. In practice, the N rows frequently correspond to the number of video frames analyzed for the given behavioral sequence. The behavior-position key will dictate which columns of the matrix correspond to which behavioral subunit. Additionally, a leading and trailing column are added (hence M = number of subunits scored + 2). The leading column will be assigned a code that is invariant across rows, as will the trailing column (though its code will be different from the leading column). These columns and the ‘start’ and ‘stop’ codes they contain are integral for aligning behavioral strings in a way that facilitates comparisons only between/among ‘like’ behavioral atom categories.

Following its creation, the behavior-time matrix is first populated by assigning 0s for every instance where a particular behavioral subunit is inactive/off and 1s for every instance when the behavioral subunit is active/on. Next, these 1s and 0s are converted into characters (or character strings) depending on the string-matching approaches to be employed. In our bird-of-paradise example (detailed below), we used behavioral sequences with a relatively limited number (31) of discrete behavioral sub-units. We substituted “0” and “1” values for each behavioral subunit with a randomly chosen, unique, triplicate nucleotide code comprised of some combination of As, Ts, Gs, and Cs. For each discrete behavioral subunit, the triplicate nucleotide code that represented the ‘on’ behavioral state was the ‘genetic complement’ of the ‘off’ behavioral state. For example, if ‘body-position-moving in space’ was active, this behavioral subunit was given the code “TTC”, whereas if this behavioral subunit was inactive the slot corresponding to this behavioral subunit was given the code “AAG”. In instances where more than 31 discrete behavioral states are used, as will commonly be the case for behavioral sequences analyzed with body-tracking tools, there are numerous alternative approaches.

When more subunits are evaluated, users can generate larger, unique sequences for each behavioral subunit state. For example, still using only A, T, G, and C, we can generate 65,536 unique sequences of 8 characters (where the first 4 characters of each sequence can be used to signify ‘on’ states, and the last 4 characters of each sequence can be used to signify ‘off’ states). Alternatively, one can generate unique character states using the protein alphabet (C, D, E, F, G, H, I, K, L, M, N, P, Q, R, S, T, V, W). For example, one can generate 18,564 unique 6 characters strings using this limited protein alphabet, again with the first 3 characters in each string corresponding to ‘on’ states, and the last 3 characters corresponding to ‘off’ states. Simply put, there are numerous ways to generate unique codes corresponding to many discrete behavioral subunits, which can then facilitate an array of bioinformatic and string-matching techniques using existing-omic toolkits.

Regardless of the specific character or character-string used to represent each behavioral subunit’s status (on/off), the last step of the RAD-behavior approach is append each row of the behavior-transition matrix, end to end, to create a single string describing the sequence of movements employed during the N time-points incorporated into the behavior-time matrix.

## Bioinformatic analyses of composite behavioral sequences

> “…many statistical techniques developed for genomic sequence analysis should be directly applicable [to the study of behavior]”^21^

Following behavioral sequence generation using the RAD-behavior framework, behavioral sequences can be numerically and quantitatively compared using string-searching and string-matching algorithms of a wide variety. To date, we have leveraged the existing computational expertise and toolkits commonly employed in the field of genomics. Specifically, by converting position-specific character states (i.e. “off” or “on”) to pseudo-genetic codes (i.e. randomly chosen nucleotide or protein sequences), we can search and compare the numerical similarity of sequences using the widely-used tool BLAST (Basic Local Alignment Search Tool). Though default settings for this program make assumptions about likelihoods and penalties of matches for different nucleotide pairs that are undoubtedly inappropriate for discretized behavioral subunits, future work collaborating with BLAST tool developers (or other computational tool developers) to eliminate the DNA-specific assumptions and calculation steps would alleviate these particular concerns. Regardless of the specific form that such future developments take, even BLAST in its current form coupled with behavioral strings generated in the RAD-behavior framework works exceedingly well at finding tight behavioral matches (i.e. sequences in which individuals are performing the same behaviors in sequence) and providing a numerical evaluation of such matches (Supplementary Video 1).

One genomic toolkit with greater flexibility, including the ability for users to define their own custom ‘alphabet’ of character states, as well the ability to implement a customized scoring matrix where users define scoring regimes is called MAFFT (Multiple Alignment using Fast Fourier Transforms)^20^. Within the RAD-behavior framework, MAFFT enables multiple behavioral sequences to be aligned and, importantly, allows for ‘insertions’ and ‘deletions’ (indels; with penalties for such discontinuities) that enables practitioners to identify “regions” of shared behavioral execution (i.e. which movement sequences are shared across individuals/contexts/etc) as well as “regions” that are unique. This approach has recently been used in an ground-breaking study examining the evolution of acoustic sequences in crickets^26^, and future investigations incorporating sequence-based analyses into phylogenetic comparative frameworks are an exciting development in the field of evolutionary biology.

In the context of behavioral strings generated via the RAD-behavior framework, multiple sequence alignment enables users to identify shared sequences of movements and, importantly, also provides the ability to highlight/identify small-scale differences in body-region-specific character states in the context of a larger sequence of movements (Figure 4). For example, MAFFT might facilitate alignment and comparison of two different individuals performing a courtship display, even if the particular cadence and tempo of the dance is different (partially via use of the insertion-deletion framework). For each shared region aligned using this framework, a user could identify which (if any) discretized behavioral states differ between the two ‘dancers’, allowing comparison of the broad-scale behavioral sequence (e.g. tail bending then extension) and fine-scale differences (e.g. ankle-bend-character X and Y shared, but ankle-bend-character Z differs).

**Figure 4.**
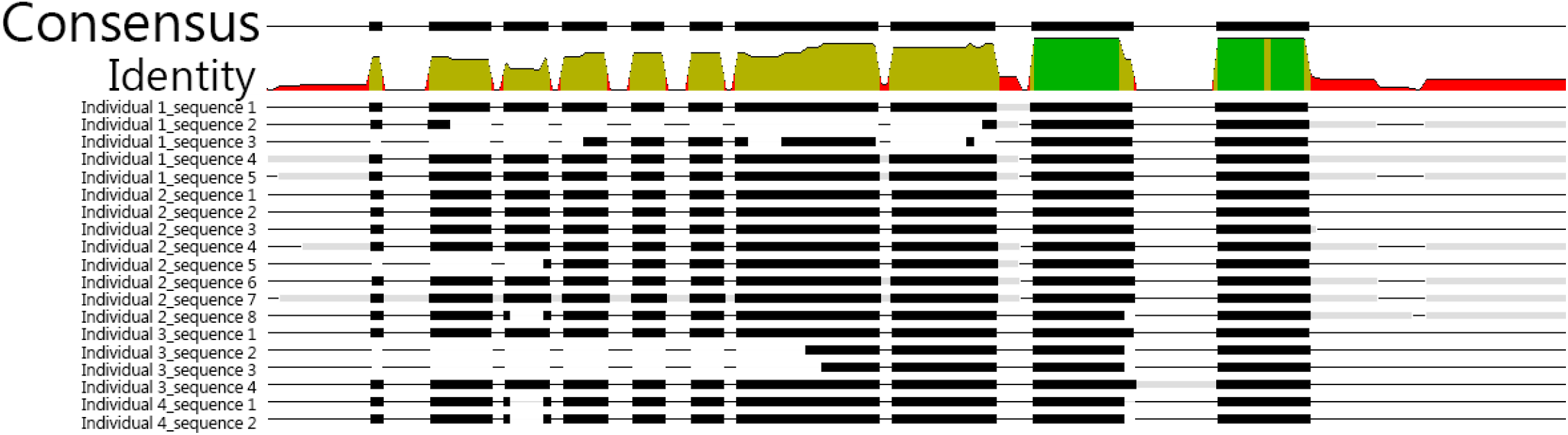
Representation of multiple sequence alignment (MSA) from four individual magnificent birds-of-paradise, each with between 2 and 8 behavioral sequences analyzed. MSA approaches highlight regions of behavioral execution that are shared with a high-degree of similarity among sequences (highlighted in green in the ‘Identity’ region of the figure), as well as those that are highly variable (low degree of consensus = red in the ‘Identity’ region of the figure).

Currently, RAD-behavior sequence comparisons have been conducted using bioinformatic tools designed to compare nucleotide and protein sequences (hence the pseudo-genetic, or pseudo-protein codes used to represent distinct behavioral subunit states, Figure 3). However, future applications of RAD-behavior would be best-suited by eliminating the genomic- and proteomic-informed assumptions in these analytical toolkits and have an extended ‘alphabet’ for coding more discretized behavioral subunits.

## Alternative implementations

In the form described above, RAD-behavior facilitates a <u>single</u> string for each behavioral sequence (e.g. courtship dance, lever pull, performance of a clinical task, etc.) describing *consecutive instantaneous configurations* of the entire (i.e. all tracked body regions) organism (e.g. how each tracked body region is positioned [e.g. where it is located, its relative orientation, its absolute orientation]) to facilitate comprehensive analyses of whole-body behavioral execution/movement.

A complementary, powerful, and case-specific usage of RAD-behavior entails the creation of <u>multiple</u> body-region-specific strings, collected simultaneously and analyzed in parallel (Figure 5). Rather than a single string describing consecutive instantaneous configurations of the entire body, this alternative approach would focus on the generation of execution strings for <u>each</u> *user-defined body region*. In this implementation, quantitative analyses and comparisons of behaviors would still be conducted on temporally-aligned behavioral sequences using string-matching/bioinformatic analyses, though output of these comparisons would necessarily be specific to the body-regions defined by the user. For example, body-region specific **reference sequences** (Box 1) could be used as a reference for comparison following the performance of a given behavioral sequence (Figure 5). In this example, multiple behavioral sequences have already been analyzed and converted to RAD-behavior execution strings, facilitating the creation of specific **consensus sequences** (Box 1) for the left wrist, left elbow, and left shoulder. Following performance of a new behavioral sequence (e.g. throwing a ball, opening a package, climbing a ladder), the new strings created for each body region can then aligned and compared to the reference sequence (in this case, a consensus sequence) facilitating rapid visualization of time-specific similarity and differences between the current execution and the consensus sequence.

**Figure 5.**
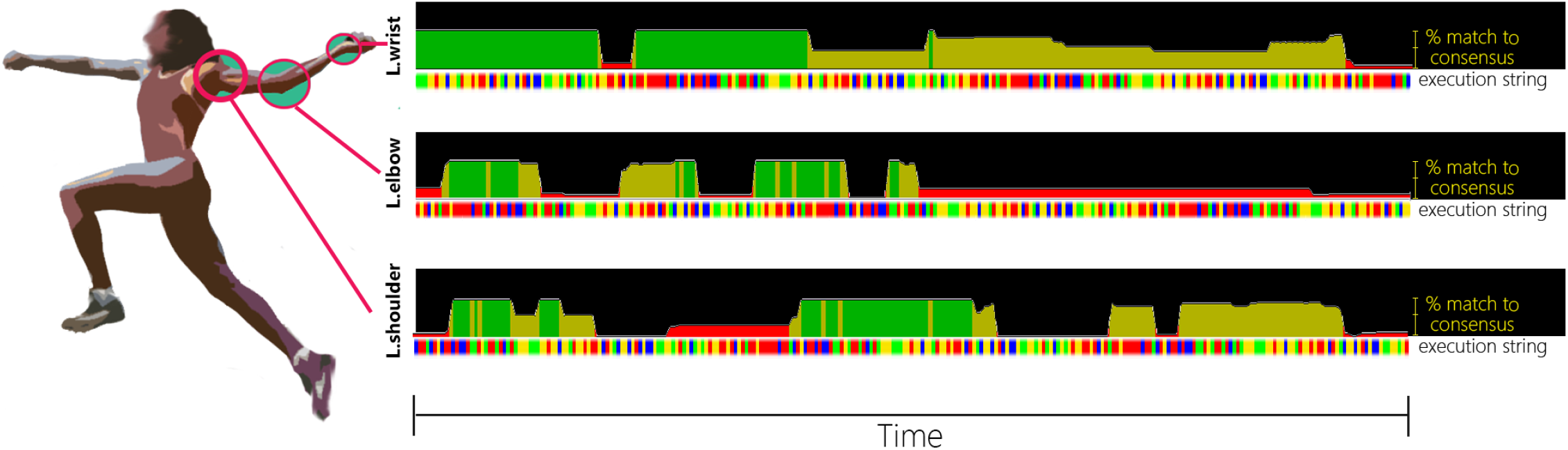
RAD-behavior sequences can be generated and analyzed for multiple user-defined body-regions independently. In this example, consensus sequences from previously analyzed behavioral sequences have been generated for the left wrist, left elbow, and left shoulder. The ‘current’ or focal behavioral execution string (with different behavioral subunits in different states indicated with different colors) is compared to this consensus string, facilitating rapid identification of body-region-specific deviations from (and matches to) the consensus (better matches equal higher % values and taller bars) sequences.

Importantly, the comprehensive and body region-specific approaches are not mutually exclusive, and the same underlying data (e.g. body-region locations in space, across time) can be used to generate both sequence sets. Users might first employ RAD-behavior to generate comprehensive strings to identify temporal periods of particular interest with respect to the overall behavioral sequence to be analyzed, and then employ the body-region-specific string generation and comparison for detailed analyses of each body part independently.

Rather than focusing on individual body-parts as the key behavioral subunits/atoms, RAD-behavior might also be used to capture and compare the behavioral execution of multiple individuals simultaneously (e.g. dyads of fighting chameleons, cooperatively hunting dolphins, courting males and evaluative females). If, rather than each body-part of the focal organism being atomized and discretized, the behavior of each team member was atomized and discretized, then combined with the discretized behavioral states of all team members, one could rapidly generate composite RAD-behavior strings detailing the movements and behaviors of the collective group across time.

## Creating rapidly searchable behavioral execution databases underlies the power of RAD-behavior

A key feature of the RAD-behavior approach to the study of behavior comes from the ability to decompose complex sequences of movements into character strings representing complex behavioral sequences which, in turn, facilitates the ability to generate (and continually add to) reference databases of behaviors that can be rapidly and efficiently searched. Behavioral databases thus enable any behavior sequence (or set of sequences) to be compared and quantitatively evaluated. This key feature is an integral component and output of the RAD-behavior pipeline. In each of the key applications described below, a reference database of behavioral sequences allows the comparisons of interest to be made.

Four broad areas of RAD-behavior application across diverse fields of application are:

A. Identifying general patterns and/or clusters of execution types (linked to particular outcomes or clustered based on execution parameters).
B. Comparing focal execution sequences to target reference sequences/databases to quantitatively assess similarity and identify where (e.g. which body regions are performing differently) and when (e.g. at what stage in a behavioral sequence, are behavioral subunits performed for different durations relative to optimal sequences) differences exist.
C. Comparing focal execution to reference databases to identify behavioral correlates (finding similarities to others based on execution of behaviors).
D. Monitoring changes in the execution of movements or sequences over time.

### A) Identification of optima or types

For many organisms, there are different ways to execute behavior sequences while still achieving individually-specific performance optima. As a human example, the world’s fastest sprinter, Usain Bolt, has a distinctive gait that would not necessarily yield ideal outcomes for another sprinter with a different body type. RAD-behavior enables execution strings to be analyzed with a series of tools that focus on patterns within long strings, similar to the approaches used in whole genome principal component analysis^31^ (PCA; Box 1). Applied to behavioral execution strings obtained with RAD-behavior, this approach would allow users to identify clusters in execution types by decomposing the high-dimensional ethomic (Box 1) data into a limited number of dimensions. For example, a user interested in geographic patterns of behavioral execution might first begin by conducting principal components analysis on a database of RAD-behavior strings. This approach would allow the user to identify which individuals/sequences/populations cluster together based on the similarity of their overall RAD-behavior derived execution strings.

### B) Comparison to optima

The factors that make a given behavioral sequence ‘optimal’ depends on the individual and species, but optimal sequences are frequently associated with some outcome connected to fitness. By connecting other data sources with RAD-behavior execution strings, we can identify ‘key’ or ‘optimal’ execution profiles (e.g. the courtship display with the greatest reproductive success in a given season), then make comparisons between the execution of non-optimal sequences and our ‘optimal’ reference sequence to see the specific ways that variation in behavioral execution might map onto traits like female choice and reproductive success. In addition to cases where we can identify a single reference sequence based on some criteria, we can also use multiple sequence alignment to generate consensus sequences (e.g. for individuals of a given age, sex, population, etc.) and evaluate individual differences from the consensus sequence on a case-by-case basis.

### C) Reference comparisons and database matches

Because the RAD-behavior approach facilitates the creation of reference databases of behavioral execution strings that can be rapidly and efficiently searched, almost any behavior (or set of behaviors) can be compared and quantitatively evaluated rapidly and efficiently. When sequences in a given database are paired with additional information about the individuals performing the behaviors (e.g. age, health-status, reproductive success, etc.), RAD-behavior string searching approaches can allow users to identify useful correlates associated with execution patterns.

### D) Longitudinal comparisons

The ability to quantitatively analyze the similarities of complex behaviors makes RAD-behavior a powerful tool to analyze changes in individual behavior over time. With RAD-behavior, numerically evaluating behavioral execution is now possible, giving the means for users to generate quantitative metrics of behavioral consistency, independent of outcome. Additionally, the bioinformatic tools that can be incorporated into a RAD-behavior analytic pipeline means users can also identify key regions where behavioral execution differs over time, thus providing new, specific, detailed insights into improvements or declines in behavioral execution (as a consequence of age, hormonal profiles, learning, etc.).

## Example demonstration using bird-of-paradise courtship behavior

Using a publicly-available database of digital media specimens (in this case, videos) located in the Macaulay Library of the Cornell Lab of Ornithology, we analyzed the courtship behavior of 31 species of birds-of-paradise from 961 video clips. Each of 31 behavioral atoms was scored manually from these clips, and the total duration of videos scored was 47,707 seconds (~13.2 hours). After repeated evaluations of each clip to facilitate fine-scale accuracy and precision in our behavioral measures, we assign behavioral states at 0.1 second intervals. Interested readers are referred to our original publication for additional details^24^ about the behavioral collection approach we employed.

Using the RAD-behavior approach, we then converted the string of consecutive, composite behavioral states obtained from each clip into a character string substituting a pseudo-genetic, triplicate nucleotide code for each behavioral atom at each time point (Supplementary Table 1). As a note, when using a pre-defined ethogram containing behavioral atoms and scoring videos manually, as we did here with the birds-of-paradise, the discretization step is implemented via the behavioral scoring of the researcher. That is, the individual who is scoring the behavior uses the description in the fine-scale, atomized ethogram to make decisions about which behaviors are ‘on’ at a given time point (with those not meeting the criteria being assigned, by default, an ‘off’ status), thus discretizing their status at every time point. Following RAD-behavior string creation, we saved the composite strings in a FASTA file (with species, clip, and individual-specific identifiers included for each substring) that contained approximately 46.6 million pseudo-nucleotides, at a file size of approximately 45 megabytes.

To illustrate the power (and limitations) of the RAD-behavior approach coupled with bioinformatic approaches, we then chose a 10 second sequence of courtship behavior from a male superb bird-of-paradise *(Lophorina superba)*, isolated the RAD-behavior sequence representing the atomized, discretized, and recombined representation of the behavioral sequence, and then BLASTed this sequence against our composite bird-of-paradise behavioral database. Following this BLAST search, we pulled the ‘best’ (i.e. highest bit-score) matches from two additional sequences containing superb bird-of-paradise courtship, and one additional match from another temporal region (i.e. non-overlapping with the query sequence) of the original recording. Interestingly, the bit-scores of sequence matching obtained in this pipeline were generally very high (Supplementary Table 2, Supplementary Video 1), likely reflecting inflated similarity scores as a consequence of inferred ‘alignment’ from sequences of both active and inactive behavioral atoms across clips/individuals. That is, much of the inferred concordance between any two sequences is as likely to come from shared ‘off’ states as shared ‘on’ states.

In agreement with the idea that the RAD-behavior approach facilitates the capture, comparison, and evaluation of behavioral sequences, the best match identified from each of three clips exhibits are large degree of overlap in many of the broad categories of behavior typically used when scoring and evaluating behavior. Specifically, all three ‘matched’ clips show the same or different male exhibiting the primary elements of courtship behavior exhibited for this species. Additionally, RAD-behavior separates the degree of concordance (and discordance) between the reference/query sequence, and the matched search strings---the top clip (blue background) demonstrates the best match in terms of overall timing and sequence similarity to the reference clip, resulting in the highest bit-score (and the lowest cumulative number of difference between it and the reference, Supplementary Figure 1, lower plot in bottom-left of Supplementary Video 1). In contrast, the third clip (bottom right, orange frame in Supplementary Video 1) shows a lower bit-score, likely reflective of the different timing of the transformation in erecting the ornamental plumage cape used during the main component of the courtship behavior of this species.

We focus on the exemplar results from BLAST in this publication for two reasons. First, the validity and power of the RAD-behavior conceptual framework can be evaluated on an intuitive level when researchers have the ability to visually evaluate and compare side-by-side those sequences determined by RAD-behavior and string-matching to be highly similar. If RAD-behavior identified sequences as being highly concordant, yet the organisms in those sequences did not look like they were performing similar behaviors when watched side-by-side, we would (rightly) be highly skeptical of proceeding with that particular framework. However, we feel that the demonstration illustrated in Supplementary Video as well as in other comparisons we have conducted but do not share here, provide strong evidence that RAD-behavior is likely to be a valuable tool aiding in the quantitative study of behavioral execution. Second, we focus on results from a RAD-behavior implementation of string-searching and matching (in the form of BLAST), because all other bioinformatic tools/approaches that we have discussed as potential future tools now useable for the study of behavior (motif identification, multiple-sequence alignment, etc.) depend on the confidence we have regarding what a string ‘match’ actually means from the perspective of behavior. Further demonstrations and implementations of RAD-behavior research on behavioral execution will test and refine these additional approaches in greater detail.

## Concluding remarks

Within the broad field devoted to behavioral tracking, identification, and comparison, RAD-behavior is unique because it generates a single string for any behavioral sequence, complex or simple. The reduction of behavior to a single string and the creation of efficiently searchable behavioral databases facilitates an array of computational comparisons of behavior that have not been possible before. Importantly, the decomposition of complex behavioral execution sequences into string format via RAD-behavior does not entail (much) loss of detailed information, as is commonly the case when complex behaviors are assigned to broader, human- or computer-defined categories, such as “running” and “eating”. Quantitative behavioral comparisons of behavioral strings obtained using RAD-behavior are also readily interpretable, in contrast to some methods that rely on dimensionality reduction. Specifically, RAD-behavior generates behavioral strings that can be quantitatively compared (analogous to the bit-scores generated when comparing genomic sequences), allowing practitioners to identify i) which behavioral sequences are most similar to one another, ii) exactly how similar sequences are (not just a ranking, but a numeric score of similarity), and iii) to readily identify, following sequence alignment, those movements/executions that are different/shared between any set of behavioral sequences (in contrast to machine-learning-based approaches which can ‘classify’ behaviors and facilitate assignment, but which do not enable practitioners to identify key differences between sequences). Further refinements, improvements, and changes to the RAD-behavior framework we have laid out here are inevitable and exceedingly welcome. Given the vast community of researchers interested in behavioral execution, there are undoubtedly a multitude of creative, innovative approaches to leverage the behavioral strings created by RAD-behavior to answer longstanding questions about behavior, as well as diverse perspectives on the challenge of identifying relevant behavioral atoms and weights. Future developments and implementations of RAD-behavior include, but are not limited to, fine-scale studies of within-individual variation in behavioral execution depending on context and learning, population-level studies focused on the role of drift and/or culture on variability in execution of shared behaviors, and interspecific comparisons of behaviors focused on homologous behavioral atoms and incorporating cutting-edge phylogenetic comparative analyses of sequence data^26^. It is our hope that RAD-behavior can serve as the conceptual framework uniting studies focused in their attempts to uncover and compare variation in behavioral evolution across scales.

---

**Supplementary Table 1.** Behavioral position-mapping key, and corresponding pseudo-genetic substitution scheme for converting bird-of-paradise courtship behavior into composite strings using RAD-behavior concepts.

| Behavior | Position | ON code | OFF code |
| --- | --- | --- | --- |
| Start | 0 | TTT |  |
| BP1 | 1 | TTC | AAG |
| BP2 | 2 | TTA | AAT |
| BP3 | 3 | TTG | AAC |
| SS1 | 4 | CTT | GAA |
| SS2 | 5 | CTC | GAG |
| O1 | 6 | CTA | GAT |
| O2 | 7 | CTG | GAC |
| O3 | 8 | ATT | TAA |
| O4 | 9 | ATC | TAG |
| O5 | 10 | ATA | TAT |
| O6 | 11 | ATG | TAC |
| O7 | 12 | GTT | CAA |
| O8 | 13 | GTC | CAG |
| OPMH | 14 | GTA | CAT |
| OPAH | 15 | GTG | CAC |
| OPABH | 16 | TCT | AGA |
| OPMB | 17 | TCC | AGG |
| OPAC1 | 18 | TCA | AGT |
| OPABB | 19 | TCG | AGC |
| OPMF | 20 | CCT | GGA |
| OPAC2 | 21 | CCC | GGG |
| OPAC3 | 22 | CCA | GGT |
| OBABF | 23 | CCG | GGC |
| OPMW | 24 | ACT | TGA |
| OPAW | 25 | ACC | TGG |
| OPATW | 26 | ACA | TGT |
| OPABW | 27 | ACG | TGC |
| OPMT | 28 | GCT | CGA |
| OPAMW | 29 | GCC | CGG |
| OPAMTT | 30 | GCA | CGT |
| OPABT | 31 | GCG | CGC |
| Stop | 31 | AAA |  |

**Supplementary Table 2.**
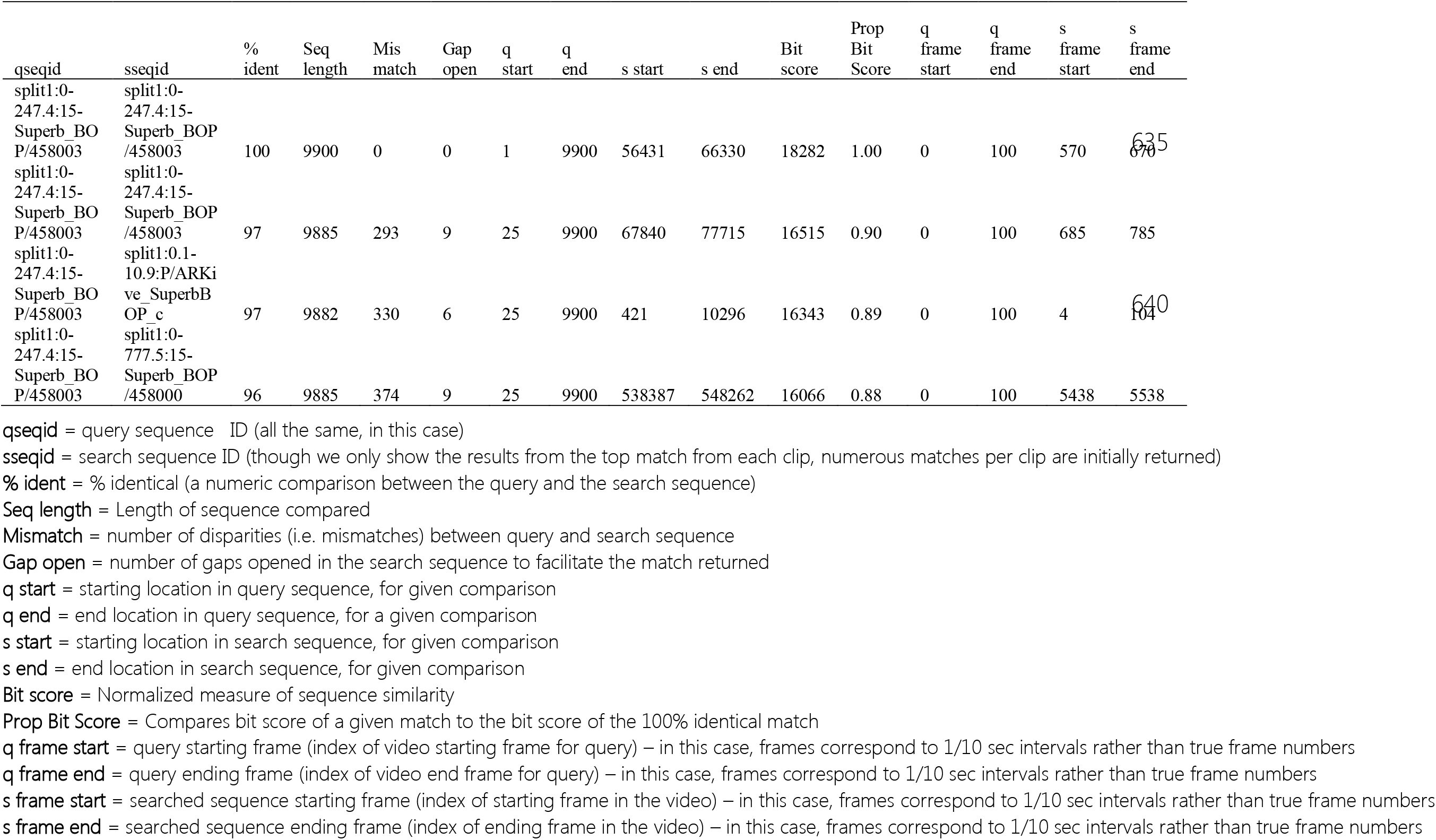
Results from a BLAST search looking for sequence matches between a reference RAD-behavior sequence (clipped from original clip <u>458003 in the Macaulay Library</u>) and database of courtship behaviors scored from 32 bird-of-paradise species (see Ligon et al. 2018^24^ for details on number of clips scored per species, as well as mean clip duration, etc.). This table (formatted based on BLAST’s tabular output, “-outfmt 6” in the command line version of BLAST), contains information about the query (i.e. reference) sequence (q) and corresponding searched (s) sequences.

**Supplementary Figure 1.**
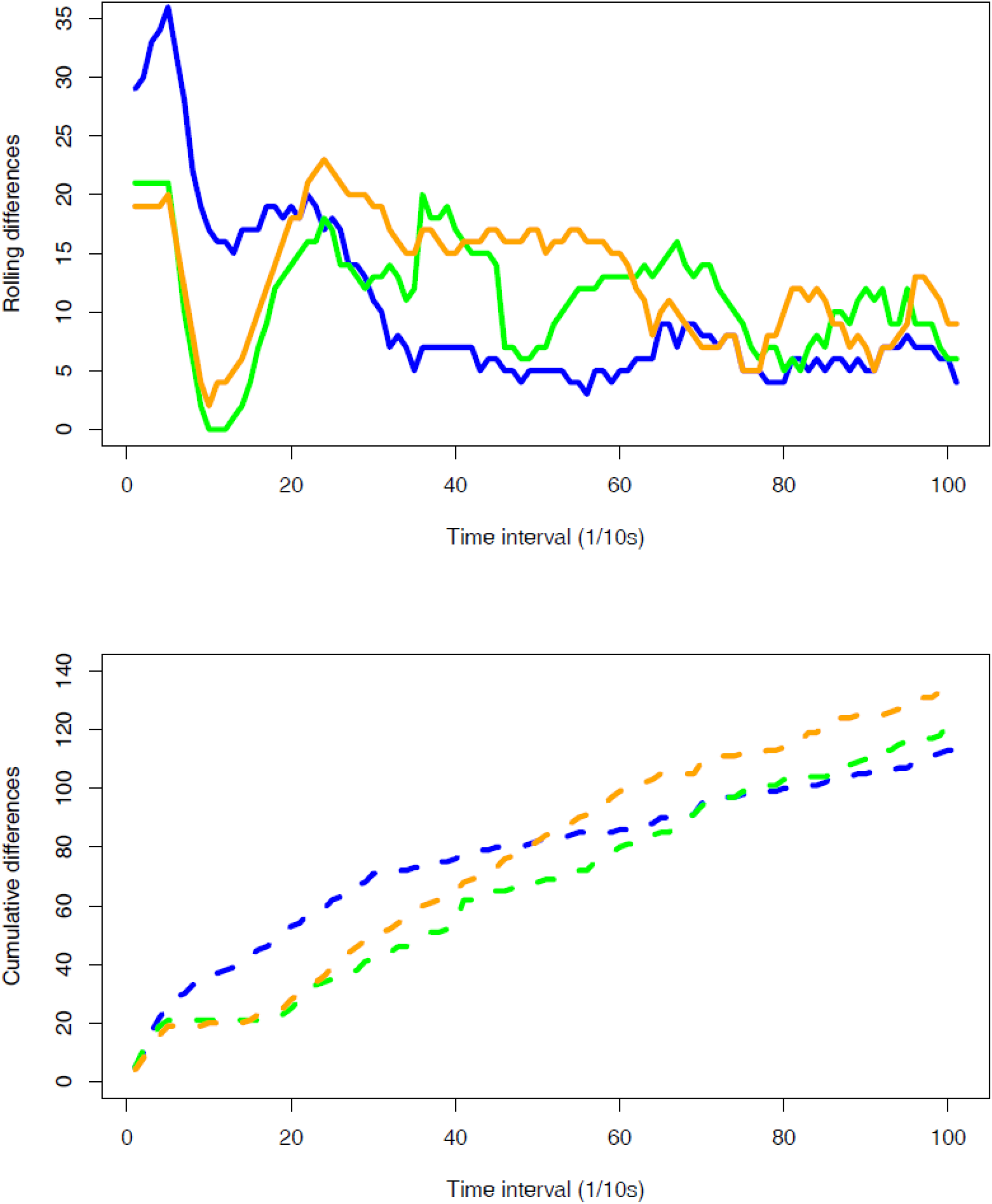
Rolling (top) and cumulative (bottom) differences between reference (query) behavioral sequence (top left of Supplementary Video 1) and three additional clips identified from a behavioral database using RAD-behavior to convert behavioral measures to character strings, then using BLAST (Base Local Alignment Search Tool) to identify sequence matches. The colors of the lines (blue, green, orange) correspond to the frame colors of the clips positioned along the right-hand side of Supplementary Video 1.

**Supplementary Video 1**. Behavior sequences scored with the RAD-behavior approach are converted to strings, which can be compiled into efficiently searchable databases. Here, a reference behavioral sequence of the courtship display of a male superb bird-of-paradise (https://macaulaylibrary.org/asset/458003, filmed by Ed Scholes III) in the top-left was searched against a cross-species database of bird-of-paradise courtship behavior using the bioinformatic tool BLAST (Basic Local Alignment Search Tool), and three of the top ‘hits’ are shown on the right side of the screen. The bit score of each clip (describing overall similarity to the reference sequence) is shown in pink (note truncated x-axis). The figure in the bottom left shows the rolling average of sequence dissimilarity between each clip (indicated by the colored boxes bounding the clips themselves) and the reference clip (top left), as well as the cumulative differences between these clips and the reference sequence. https://youtu.be/naAySqzVnNo

## Literature Cited

1. Gernat T, Rao VD, Middendorf M, Dankowicz H, Goldenfeld N, Robinson GE, Holme P, Naug D, Sokolowski MB. 2017. Automated monitoring of behavior reveals bursty interaction patterns and rapid spreading dynamics in honeybee social networks. 1–6 doi:10.1073/pnas.1713568115

2. Mathis A, Mamidanna P, Abe T, Cury KM, Murthy VN, Mathis MW, Bethge M. 2018. Markerless tracking of user-defined features with deep learning. 1–14

3. Aoki R, Tsubota T, Goya Y, Benucci A. 2017. An automated platform for high-throughput mouse behavior and physiology with voluntary head-fixation. Nat. Commun. 8: 1196

4. Geissmann Q, Garcia Rodriguez L, Beckwith EJ, French AS, Jamasb AR, Gilestro GF. 2017. Ethoscopes: An open platform for high-throughput ethomics. PLoS Biol. 15: 1–13

5. Mersch DP, Crespi a., Keller L. 2013. Tracking Individuals Shows Spatial Fidelity Is a Key Regulator of Ant Social Organization. Science (80-.) 340: 1090–1093

6. Branson K, Robie AA, Bender J, Perona P, Dickinson MH. 2009. High-throughput ethomics in large groups of Drosophila. Nat. Methods. 6: 451–457

7. Dankert H, Wang L, Hoopfer ED, Anderson DJ, Perona P. 2009. Automated monitoring and analysis of social behavior in Drosophila. Nat. Methods. 6: 297–303

8. Gomez-Marin A, Paton JJ, Kampff AR, Costa RM, Mainen ZF. 2014. Big behavioral data: psychology, ethology and the foundations of neuroscience. Nat. Neurosci. 17: 1455–1462

9. Brown AEX, de Bivort B. 2018. Ethology as a physical science. Nat. Phys. 1–5 doi:10.1038/s41567-018-0093-0

10. Berman GJ. 2018. Measuring behavior across scales. BMC Biol. 16: 23

11. Dominici N, Ivanenko YP, Cappellini G, Avella A, Mondì V, Cicchese M, Fabiano A, Silei T, Paolo A Di, Giannini C, Poppele RE, Lacquaniti F. 2011. Locomotor Primitives in Newborn Babies and Their Development. 334: 997–1000

12. Whitman CO. 1898. in *Biol Lect. from Mar. Biol Lab*. wood’s holl[sic]. 285–338 Ginn & Co.

13. Heinroth O, Heinroth M. 1928. Die Vogel Mitteleuropas. Bermuhler

14. Lorenz KZ. 1941. Vergleichende Bewegungsstudien an Anatinen. J. für Ornithol 79: 194–294

15. Dewsbury DA. 1978. Comparative Animal Behavior. McGraw-Hill

16. Eshkol N, Wachman A. 1958. Movement Notation. Weidenfeld & Nicolson

17. Hutchinson A. 1954. Labanotation. Routledge

18. Altschul SF, Gish W, Miller W, Myers EW, Lipman DJ. 1990. Basic Local Alignment Search Tool. J. Mol. Biol. 215: 403–410

19. Bailey TL, Boden M, Buske FA, Frith M, Grant CE, Clementi L, Ren J, Li WW, Noble WS. 2009. MEME SUITE: tools for motif discovery and searching. Nucleic Acids Res. 37: W202–W208

20. Katoh K, Standley DM. 2013. MAFFT multiple sequence alignment software version 7: improvements in performance and usability. Mol Bioil. Evol. 4: 772–780

21. Reiser M. 2009. The ethomics era? Nat. Methods. 6: 413–414

22. Nissen HW. 1958. in Behav Evol. Roe A, & Simpson GG, ed. 183–205 Yale University Press

23. Berman GJ, Bialek W, Shaevitz JW. 2016. Predictability and hierarchy in Drosophila behavior. 1–6 doi:10.1073/pnas.1607601113

24. Ligon RA, Diaz CD, Morano JL, Troscianko J, Stevens M, Moskeland A, Laman TG, Scholes E. 2018. Evolution of correlated complexity in the radically different courtship signals of birds-of-paradise. PLoS Biol. 16: e2006962

25. Gkoutos G V, Hoehndorf R, Tsaprouni L, Schofield PN. 2015. Best behaviour? Ontologies and the formal description of animal behaviour. Mamm. Genome. 26: 540–547

26. Caetano DS, Beaulieu JM. 2019. Comparative analyses of phenotypic sequences using phylogenetic trees. bioRxiv http://dx.doi.org/10.1101/561167

27. Jackson BE, Evangelista DJ, Ray DD, Hedrick TL. 2016. 3D for the people: multi-camera motion capture in the field with consumer-grade cameras and open source software. J. Exp. Biol. 5: 1334–1342

28. Berman GJ, Choi DM, Bialek W, Shaevitz JW. 2014. Mapping the stereotyped behaviour of freely moving fruit flies. J R Soc Interface. 11: 20140672

29. Mathis A, Mamidanna P, Cury KM, Abe T, Murthy VN, Mathis MW, Bethge M. 2018. DeepLabCut: markerless pose estimation of user-defined body parts with deep learning. Nat. Neurosci. doi.org/10.1038/s41593-018-0209-y

30. Pereira TD, Aldarondo DE, Willmore L, Kislin M, Wang SS, Murthy M, Shaevitz JW. 2019. Fast animal pose estimation using deep neural networks. Nat. Methods. 16: 117–125

31. Abraham G, Inouye M. 2014. Fast Principal Component Analysis of Large-Scale Genome-Wide Data. 9: 1–5

